# SATB2 loss in inflammatory bowel disease-associated small intestinal metaplasia of the distal colon

**DOI:** 10.1101/2023.02.01.526729

**Authors:** Maged Zeineldin, Tatianna C. Larman

**Author notes:** Correspondence: Tatianna Larman, MD, 855 N Wolfe St, Rangos 453, Baltimore, MD 21205.

## Abstract

Epithelial metaplasia is a common adaptation to chronic inflammatory processes and can be associated with increased risk of dysplasia and cancer. The distal colon of patients with inflammatory bowel disease (IBD) commonly shows crypt architectural distortion and Paneth cell metaplasia (PCM), and IBD patients also carry increased risk of colitis-associated dysplasia and cancer (CAC). Loss of SATB2 expression (Special AT-rich binding 2 protein, a colon-restricted chromatin remodeler) has recently been shown to distinguish colitis-associated dysplasia and CAC from sporadic disease. Here we report non-diffuse heterogeneous patterns of SATB2 loss across non-dysplastic distal colon biopsies from IBD patients (n=20). This cohort was specifically curated to include biopsies with well-developed histologic features of villiform growth and PCM. Notably, CDX2 was strongly expressed and P53 showed a wild-type immunolabeling pattern across our non-dysplastic cohort, regardless of SATB2 immunolabeling pattern. Our findings fit with recent murine studies in which colon-specific *Satb2* deletion resulted in histologic conversion of colonic mucosa to small intestinal-like mucosa, including emergence of villi and Paneth cells. Taken together, we show that SATB2 loss is associated with a preneoplastic metaplastic response to chronic injury in human IBD and chronic colitis, reframing PCM more broadly as small intestinal metaplasia. We propose that inflammation-associated SATB2 loss mediates a remodeled chromatin landscape permissive for dysplasia and CAC.

## Introduction

Metaplasia, or the histologically evident replacement of one differentiated somatic cell type with another, offers insights into mucosal homeostasis as an adaptation to diverse inflammatory pathogenic stimuli^1^. Tissue identity is dictated by interactions between tissue-specific transcription factors (TFs) and epigenetic enhancers which lead to lineage-specific transcriptional programs^2^. Inflammation can change the expression of TFs, epigenetic regulators, and/or chromatin remodelers, altering cell identity and resulting in metaplasia^2,3^. Inflammation-associated metaplasia is commonly associated with increased risk of epithelial dysplasia and cancer^1^.

Inflammatory bowel disease (IBD) is an etiologically complex chronic inflammatory disease characterized by relapsing cycles of intestinal injury and healing^4^. IBD patients carry increased risk of colitis-associated dysplasia and colorectal cancer (CAC) proportional to disease duration, extent, and severity^5^. Chronic inflammation in IBD is also associated with so-called Paneth cell metaplasia (PCM) in the distal colon as well as crypts with architectural distortion and villiform growth^6^. The molecular mediators of this epithelial and mucosal remodeling in chronic IBD are not known^7–9^.

SATB2, or special AT-rich binding protein 2, is a chromatin remodeling protein with homeostatic roles in osteoblastic and neural differentiation^10^. Importantly, SATB2 is also a specific and robust marker of colon intestinal identity^11^. CDX2, by contrast, is a TF marker of both small intestinal and colonic differentiation^11^. The critical role of SATB2 in orchestrating colonic epithelial identity was recently revealed when *Satb2* deletion in murine colonic epithelium resulted in striking conversion of colonic mucosa to small intestine, including emergence of villi and Paneth cells^12^. Prior work has also shown that bone morphogenic protein-directed SATB2 expression is important in defining colonic identity in human colon organoids derived from induced pluripotent stem cells^13^.

Intriguingly, SATB2 expression (at the protein level) is uniquely lost in human IBD dysplasia and CAC, in contrast to sporadic CRC where its expression is typically retained^14,15^. SATB2 loss has also recently been associated with changes indefinite for dysplasia and risk of subsequent dysplasia^16^. By contrast, CDX2 expression is retained in IBD-associated neoplasia^14,15^. Notably, while mutations in the tumor suppressor gene *TP53* are common early events in IBD dysplasia and CAC, genetic alterations in *SATB2* have not been reported^17,18^.

Although prior studies^14–16^ showed that SATB2 expression was reportedly overall intact in non-dysplastic active IBD biopsies, we wondered whether SATB2 loss could be seen specifically in the specific histologic context of distal chronic colitis with features of “small intestinalization,” including PCM and villiform distorted growth. Here we show that SATB2 loss is indeed common in this “small intestinal metaplasia” histologic phenotype and propose that SATB2 plays a role in a metaplasia-dysplasia sequence in IBD.

## Materials and methods

Colon samples distal to the splenic flexure were selected from patients with established history of IBD (n=6 Crohn’s disease; n=14 ulcerative colitis) from the Johns Hopkins Hospital formalin fixed paraffin embedded tissue archives under an IRB-approved protocol. H&E stains were reviewed by an expert gastrointestinal pathologist (T.L.) to identify non-dysplastic biopsies with well-developed crypt architectural distortion and PCM. Patients with known history of colitis-associated dysplasia and CAC were excluded. Our cohort additionally included 3 cases of non-IBD chronic colitis (diverticular-associated disease); normal terminal ileum (n=3); colitis-associated low-grade dysplasia (n=2); colitis-associated high-grade dysplasia (n=1); and colitis-associated cancer (n=1).

Immunostaining was performed with clinically-validated antibodies on unstained tissue sections as follows: SATB2 (clone EP281, Cell Marque, Rocklin, CA), p53 (clone Bp53-11, Roche, Indianapolis, IN), and CDX2 (DAK-CDX2, Agilent/DAKO, Santa Clara, CA). SATB2 immunostaining was interpreted as lost/attenuated if epithelium in crypts showed less expression compared to internal positive controls as evaluated by brightfield microscopy.

## Results

### SATB2 epithelial expression is frequently attenuated in non-dysplastic distal IBD colon with architectural distortion and Paneth cell metaplasia

We first verified expected SATB2 expression patterns in normal terminal ileum, normal colon, colitis-associated dysplasia, and CAC. As reported previously^12,13^, normal terminal ileum shows negative to a faint blush of nuclear staining, whereas normal colon shows diffuse nuclear expression more pronounced in crypt bases (**Fig 1**). As expected^15,16^, all three cases of colitis-associated dysplasia and CAC showed diffuse loss of nuclear SATB2 immunolabeling in foci of dysplasia and CAC, with intact CDX2 expression and mutant-pattern of P53 immunolabeling (**Fig 1**). SATB2 expression was intact in surrounding non-dysplastic epithelium, as previously reported^14,15^.

**Figure 1.**
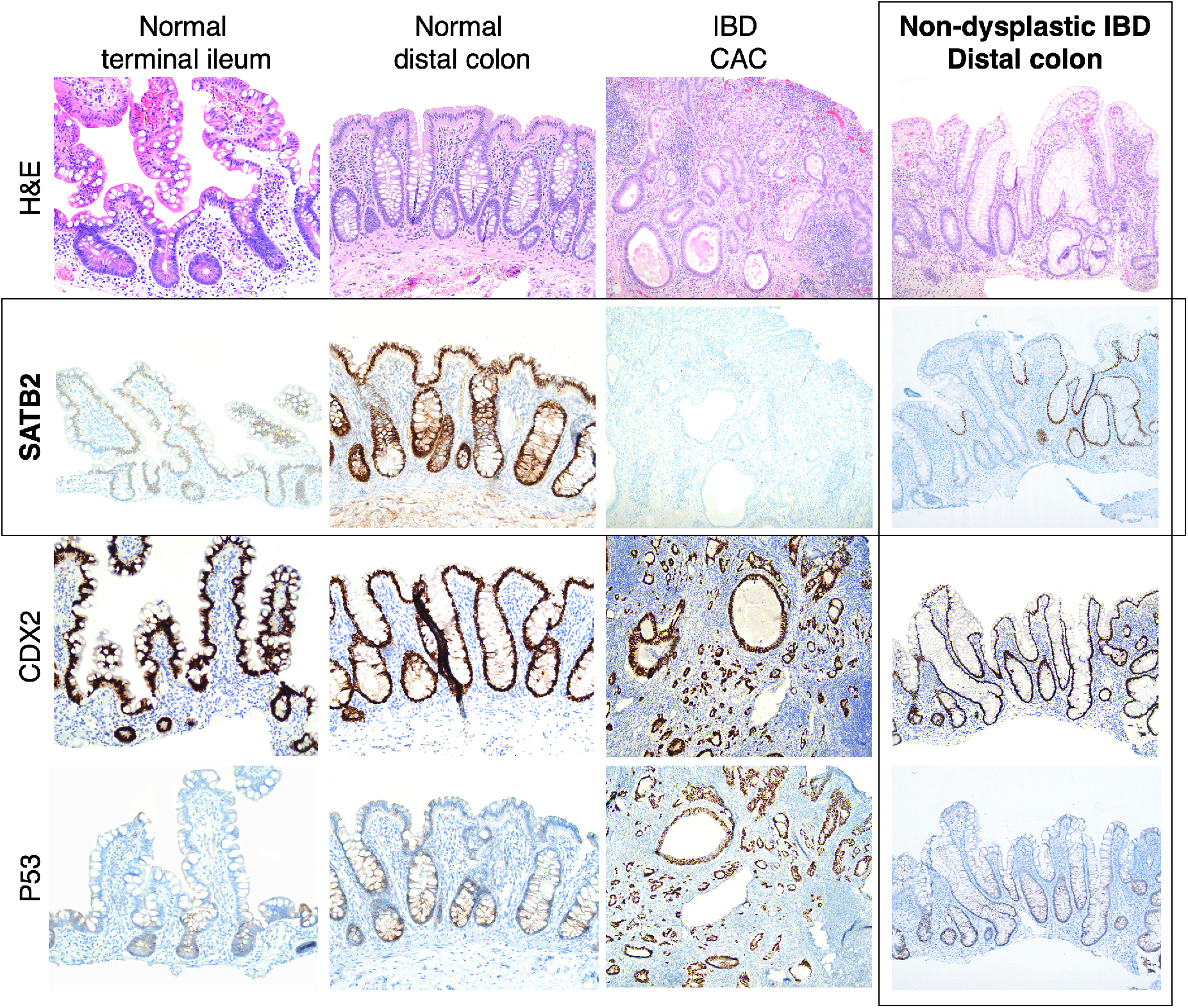
Loss of SATB2 protein expression is commonly seen in the setting of chronic IBD involving distal colon with architectural distortion and Paneth cell metaplasia. Representative H&E images and immunolabeling from each cohort.

Strikingly, all distal colon chronic IBD biopsies in our series showed some degree of SATB2 attenuation in crypts. 11 out of the 20 (55%) showed foci of complete loss involving entire or contiguous crypts (**Fig 1, Fig 2A**). These patterns of SATB2 loss were seen in both Crohn’s disease and ulcerative colitis. Remaining biopsies showed more focal/patchy patterns of SATB2 attenuation, even within the same crypt. As expected, CDX2 immunolabeling (performed in a subset of non-dysplastic samples, n=6), stained all epithelial cells with uniform intensity regardless of SATB2 immunolabeling status. P53 accordingly showed a wild-type immunolabeling pattern (performed in a subset of non-dysplastic samples, n=6) (**Fig 1**).

**Figure 2.**
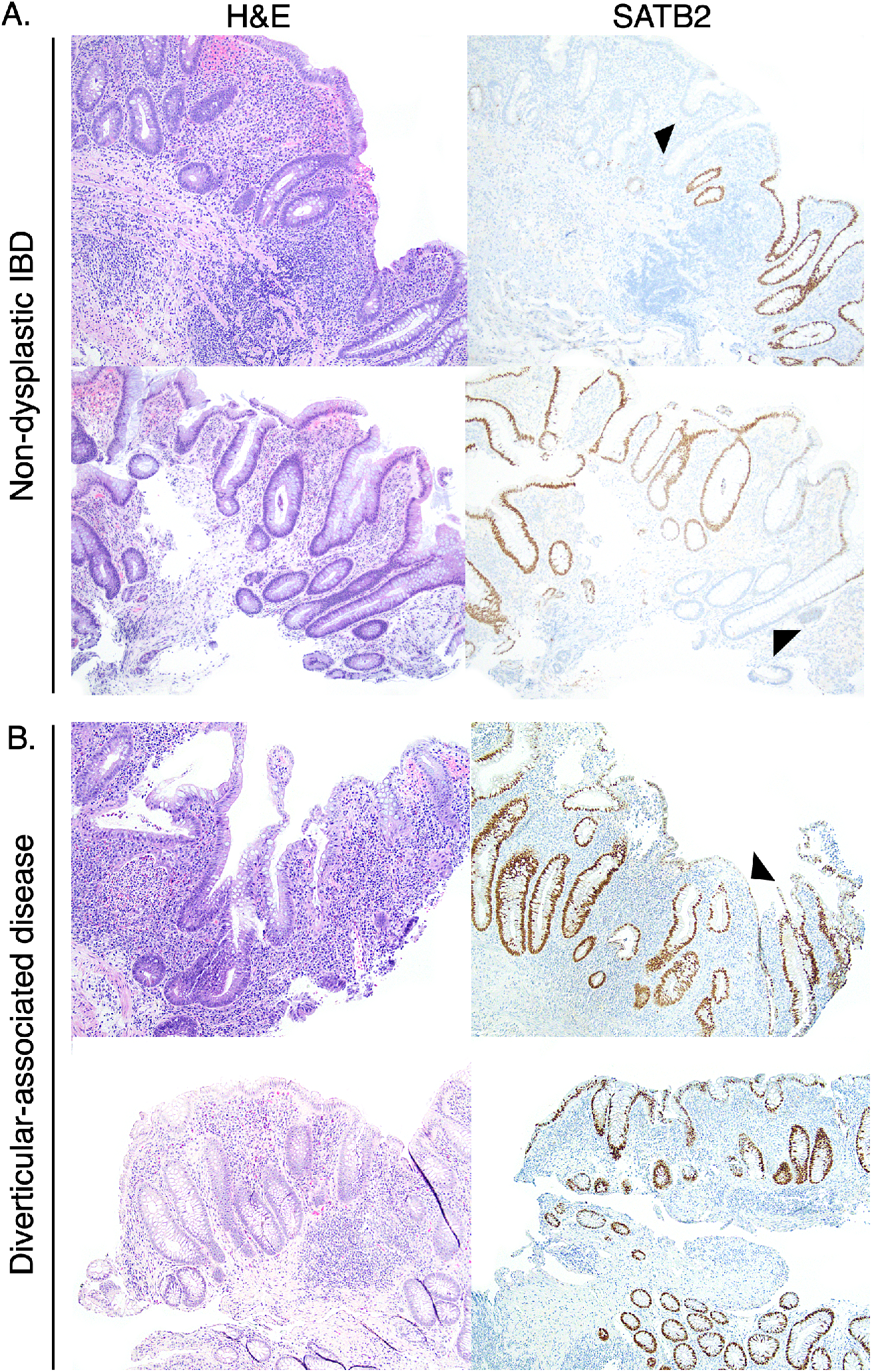
Comparison of patterns of SATB2 loss in human IBD (A) vs. non-IBD active chronic colitis involving the distal colon (B). Arrowheads denote foci of lost/attenuated SATB2 immunolabeling. SATB2 loss more commonly involved contiguous crypts in IBD (A) whereas SATB2 immunolabeling exhibited only focal attenuation (upper panel) or overall intact expression (lower panel) in non-IBD chronic colitis (B).

### SATB2 loss is observed in non-IBD chronic colitis

Histologic features of chronic injury (PCM, crypt architectural distortion) seen in the distal colonic mucosa of IBD are not specific to IBD^19^. We wondered whether SATB2 loss is a molecular change specific to IBD-associated chronic injury or whether it can also be seen in non-IBD forms of chronic colitis. To address this, our cohort included 3 distal colon biopsies from patients with diverticular-associated disease with PCM and architectural distortion. Two cases showed foci of SATB2 attenuation, although not to the degree seen in IBD. One case showed intact SATB2 expression (**Fig 2B**).

## Discussion

Here we link SATB2 loss to a chronic mucosal injury phenotype in human IBD. This study contributes to the discussion of SATB2 in IBD dysplasia and its role in normal intestinal homeostasis.

To study the role of SATB2 in IBD metaplasia/dysplasia, we included only distal colon biopsies with well-developed chronic features of IBD (marked crypt distortion and PCM), regardless of active inflammation. The specificity of our inclusion criteria may explain the apparent discordance of our results and previous studies, which reported overall intact SATB2 expression in the majority of non-dysplastic active IBD biopsies^14–16^. However, these previous studies profiled active IBD biopsies from any intestinal segment. As the composition of our cohorts differ in a biologically relevant way, we believe our findings are overall compatible with and contextualize this prior work.

Our data reveal subtleties to SATB2 loss patterns that raise the possibility that SATB2 loss in non-dysplastic IBD could be a reversible and dynamic phenomenon. For example, foci of PCM are not always correlated with foci of SATB2 loss (**Fig 2A**). Several potential (non-exclusive) possibilities could explain these phenomena. SATB2 expression could be lost and re-gained upon resolution of injury, and our biopsies capture a moment in time along this spectrum. Histologic findings could also represent proximal colon reprogramming/differentiation (as proximal colon normally has Paneth cells and SATB2 expression) of unknown etiology, rather than truly small intestinal reprogramming. Finally, other unknown molecular could mediate PCM and small intestinalization.

Metaplasia is considered an adaptive response to pathologic inflammation^1^. This raises the question of how small intestinal metaplasia could be advantageous for injured colonic epithelium. There is rich literature of Paneth cells promoting and establishing an intestinal stem cell niche^20,21^, and potentially PCM could promote a regenerative response to injury. Recent work has also implicated increased autophagic activity (seen in Paneth cells) as promoting regeneration and epithelial resilience in injury^22–24^.

Limitations of our study include that we did not correlate SATB2 expression with IBD disease duration or treatment modality. In addition, the size of our non-IBD chronic colitis cohorts is relatively limited. As the aim of the study was to examine SATB2 expression in IBD-associated PCM (seen only in the distal colon), we did not assess for SATB2 loss in IBD involving the proximal colon.

In sum, we report SATB2 loss as a phenomenon commonly seen in the distal colon of chronic IBD (and possibly other forms of chronic colitis) that contributes to a small intestinal metaplasia histologic phenotype. Due to the role of SATB2 in chromatin remodeling^12^, we posit that modulation of SATB2 expression in IBD is a potential non-genetic mechanism for a remodeled chromatin landscape permissive for neoplasia, aneuploidy, and *TP53* alterations^17,18^. Future work will investigate mediators and mechanisms of inflammation-associated SATB2 loss. Other future directions include investigation of how “small intestinalization” of colon could locally perpetuate chronic inflammatory disease, for example by impairing barrier function or promoting dysregulated immune responses and/or microbial dysbiosis.

